# Segmenting nuclei in brightfield images with neural networks

**DOI:** 10.1101/764894

**Authors:** Dmytro Fishman, Sten-Oliver Salumaa, Daniel Majoral, Samantha Peel, Jan Wildenhain, Alexander Schreiner, Kaupo Palo, Leopold Parts

## Abstract

Identifying nuclei is a standard first step to analysing cells in microscopy images. The traditional approach relies on signal from a DNA stain, or fluorescent transgene expression localised to the nucleus. However, imaging techniques that do not use fluorescence can also carry useful information. Here, we demonstrate that it is possible to accurately segment nuclei directly from brightfield images using deep learning. We confirmed that three convolutional neural network architectures can be adapted for this task, with U-Net achieving the best overall performance, Mask R-CNN providing an additional benefit of instance segmentation, and DeepCell proving too slow for practical application. We found that accurate segmentation is possible using as few as 16 training images and that models trained on images from similar cell lines can extrapolate well. Acquiring data from multiple focal planes further helps distinguish nuclei in the samples. Overall, our work liberates a fluorescence channel reserved for nuclear staining, thus providing more information from the specimen, and reducing reagents and time required for preparing imaging experiments.

## Introduction

Much of our understanding of cells has been derived from microscopy experiments. Images informed the very definition of the word “cell” (Hooke and Jo Martyn And 1665) to describe their structure, led to the discovery of organelles like nuclei (Brown 1833) and mitochondria (Altmann 1894), and are currently central to phenotyping individual cells (Usaj et al. 2016). Advances in instrumentation have enabled scalable approaches to visualize and quantify cell features, which are widely used in basic research and drug development (Li et al. 2016).

High throughput fluorescence cell imaging collects information from a small number of partially overlapping emission spectra. Each of these can correspond to the expression of a protein to reflect its abundance and localization in the cell or a stain that is enriched in areas with particular biochemical properties. The former are typically used to study individual genes, while the latter usually mark entire organelles. Popular staining choices are Hoechst dyes for DNA (and thereby, cell nuclei), mitotracker for mitochondria, phalloidin for actin, concanavalin A for lectins, and wheat germ agglutinin for membranes (Bray et al. 2016). The formed structures are easily visually distinguished and can be accurately detected with automated computational methods (Rohban et al. 2017; Pärnamaa and Parts 2017).

While cell stains are useful, there are drawbacks to using them. Only a limited number of non-overlapping stains can be used on one specimen. This is a bottleneck in experiments where stains are used to record the layout of organelles as a reference for measuring individual proteins, as each organelle takes up a channel that could be used to study a gene. The process of staining itself takes substantial time and resources. If cells need to be fixed, it destroys the sample and therefore precludes recording dynamic information, such as response to a drug (Isherwood et al. 2011). It is also possible to stain live cells, but such reagents tend to be reactive and leak from the organelles, making interpretation of other effects difficult. Common live imaging alternatives are cell line constructs with fluorescent labeling that either need to be purchased or laboriously engineered. Therefore, reducing the number of objects that need to be labeled is of great practical importance.

Brightfield images are a source of complementary information about a sample. This imaging modality records natural light transmission properties, and is therefore not specific to any particular structure. As nuclei contain densely packed DNA, photons passing through them take an optical path different from the surrounding cytoplasm, which can in principle be used to identify their location. If feasible, using brightfield images for detecting nuclei would enable experiments with an additional free channel for staining, or possibility for the temporal acquisition of live cells without perturbing their state.

The main barrier for widespread use of brightfield images for nuclear segmentation is the difficulty of automated image analysis. This is a complicated task even for humans, and as classical computational approaches have not been sufficiently accurate, fluorescent staining with its clean nuclear signal has been preferred. However, recent breakthroughs in deep learning have led to impressive performance on image analysis tasks in general (Angermueller et al. 2016; Fan and Zhou 2016), and segmentation from cell images in particular (Van Valen et al. 2016; Falk et al. 2019; Christiansen et al. 2018; Jones et al. 2017). This motivates a re-evaluation of whether nuclear segmentation could be achieved without a DNA stain.

Here, we investigate whether deep learning methods can help segment nuclei in brightfield images. We compare the performance of three convolutional neural network architectures that have been previously used for segmentation tasks, and apply the best performing one on a panel of images from seven different cell lines for which both fluorescence and brightfield images have been acquired. We evaluate how much data is required to train a successful network, and whether multiple focal planes aid segmentation.

## Results

To evaluate models, we used a hand-curated gold standard dataset of seven cell lines for which both fluorescent readout of a DNA-binding dye, and a brightfield measurement were acquired (Figure 1A). The ground truth nuclear segmentations were generated from the fluorescence channel with the PerkinElmer Harmony software with the manual tuning of the parameters and expert quality control. We considered three popular neural network architectures for image analysis -- DeepCell (Van Valen et al. 2016), U-Net (Falk et al. 2019), and Mask R-CNN (He et al. 2018), and performed a further architecture search to select the best model within each class according to performance on a held-out test set (Methods).

**Figure 1.**
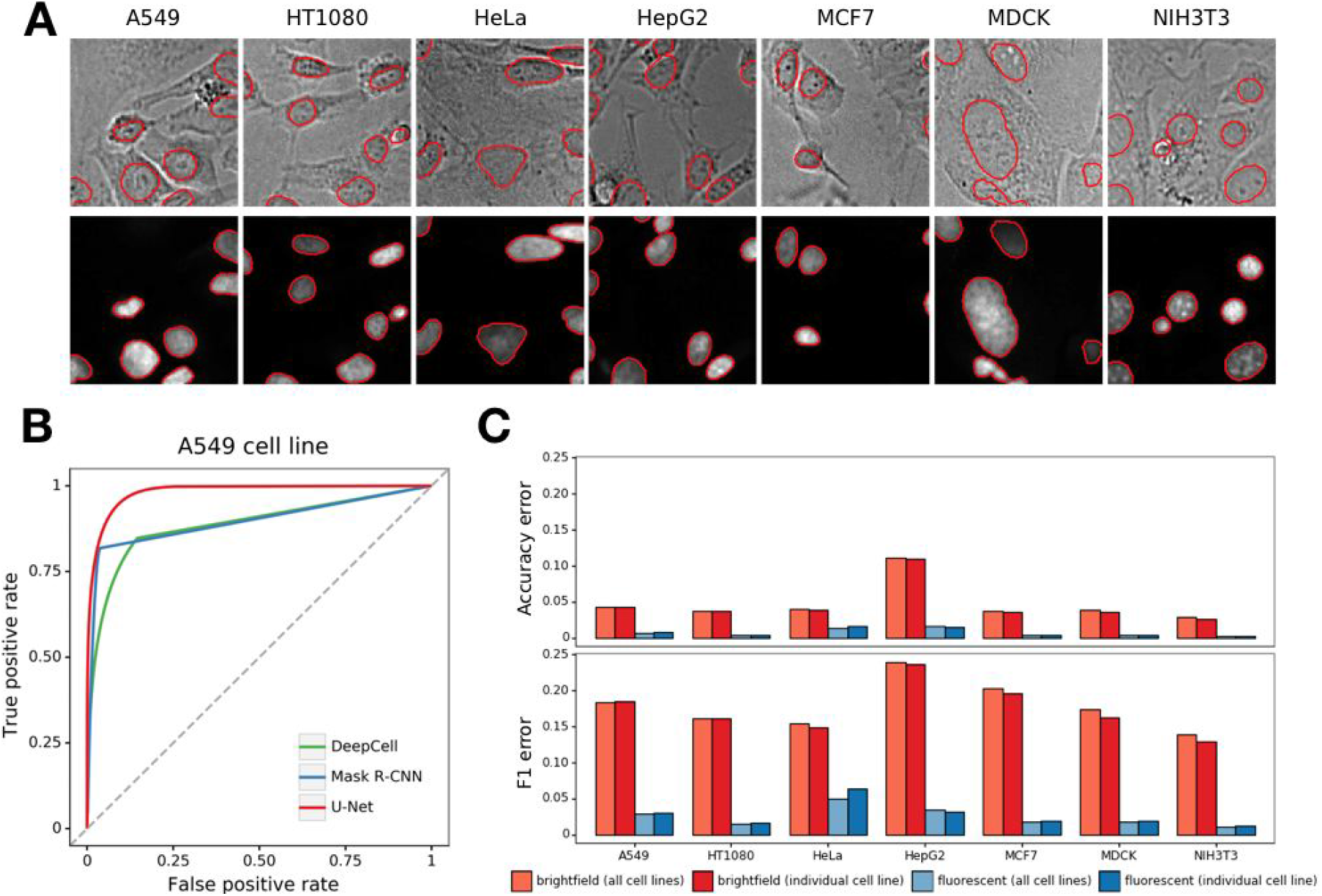
A combined fluorescence and brightfield dataset for nuclear detection. **A**. Microscopy images were acquired from seven cell lines (x-axis) using the brightfield channel (top) and green fluorescence (bottom); a representative patch from a larger image is shown for each. The ground truth nuclear boundary from the fluorescence channel is overlaid in red. Segmentation was performed in PerkinElmer Harmony software with manual parameter tuning. **B**. Neural networks successfully segment nuclei from brightfield images. Pixel-level true positive rate (y-axis) and false positive rate (x-axis) at different score cutoffs for U-Net (red), DeepCell (green), and Mask R-CNN (blue). Grey dashed line: *y*=*x*. **C**. U-Net segmentation error (y-axis) for different cell lines (x-axis) using unbalanced (1 – accuracy, top) and balanced (1 – F1 score, bottom) error measures for brightfield (red) and fluorescence (blue) channels, and models trained on one cell line at a time (light) as well as on all data (dark).

### Deep learning models can effectively segment brightfield images of different cell lines

We first evaluated the selected models on brightfield images from the A549 cell line. While this data modality is visually challenging, all approaches performed better than a random classifier (Figure 1B). U-Net achieved the best accuracy overall, with an area under receiver operating characteristic (AUROC) of 0.98, and accuracy of 96%. Mask R-CNN had a lower AUROC of 0.89, but similar accuracy of 96%, while DeepCell was less accurate (90%, 0.89 AUROC). The performance difference was consistent across test images, with U-Net always more accurate than Mask R-CNN, and the latter more accurate than DeepCell (Figure S1).

We next evaluated the U-Net model further on all cell lines, and both the fluorescence and brightfield channel. Brightfield channel performance varied across cell lines (accuracy 0.89 to 0.97, F1-score 0.76 to 0.86), while fluorescent images were consistently well segmented (accuracy 0.98 to 0.998 and F1 score 0.95 to 0.99, Figure 1C). This bias in performance between data modalities is not surprising, as the ground truth is derived from fluorescence readout. HepG2 cells were the most difficult to segment in the brightfield channel, likely due to the high cell density in images, which was negatively correlated with per-image accuracy (Figure S2). Altogether, nuclei can be consistently segmented from both brightfield and fluorescence images regardless of the cell line.

We next trained the best performing U-Net architecture on images from each cell line and channel separately. The individual line models had 0.5% lower error compared to the network trained on all data (17.4% to 17.9% average error, Figure 1C; Table S2, S3) for the brightfield channel. Fluorescent channel segmentation error increased slightly in the individual cell line models, but the difference was small in absolute terms (0.05% on average; Table S4, S5). Overall, while performance varies when only a few training images are available, there is little performance difference between models trained with ample data on the target cell line only, and ones trained on a broader training set.

### Segmentation depends on the network architecture

The alternative network architectures lead to qualitatively different segmentation outcomes. U-Net uses a model where the posterior probabilities of nearby pixels are coupled, resulting in a smooth probability map across the image (Figure 2A), and allowing trading sensitivity for specificity while maintaining overall object shape. In contrast, DeepCell classifies individual pixels based on a small surrounding area, resulting in unclear boundaries, and speck noise (Figure 2B). Finally, Mask R-CNN considers proposal regions, which are accepted or rejected as a whole, and the accepted regions further binary segmented. This results in sharp decision boundaries, and entire regions collectively assigned to foreground or background (Figure 2C).

**Figure 2.**
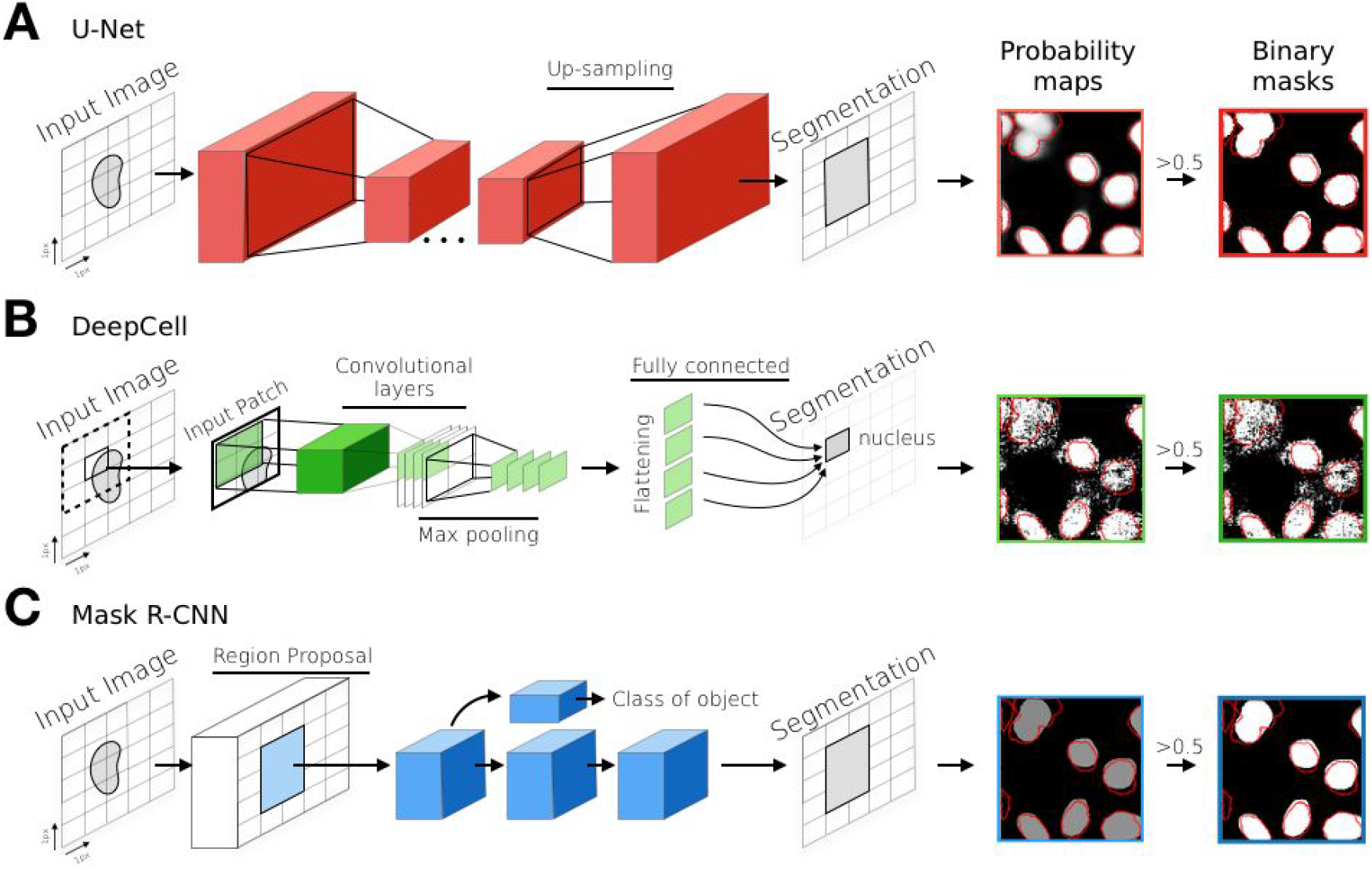
Deep learning models for nuclear segmentation. **A**. U-Net goes through a series of down- and then up-sampling steps. **B**. DeepCell considers a single pixel in its context at a time. **C**. Mask R-CNN combines multiple networks for proposing regions, classifying them, and segmenting foreground pixels.

We inspected the outputs of all test images for the U-Net model, and made the following observations about various types of mistakes in segmentation. The recall of ground truth pixels for each cell was bimodal, with some cells completely missed, and others with incorrect boundaries (Figure S3). Some nuclei were missed completely, either due to having been filtered out when generating ground truth, appearing in cells with unusual morphology (Figure S4), or having very low signal (Figure S5). Nuclei densely packed in the fluorescent channel, likely often reflecting partially overlapping cells, were more difficult to delineate correctly (Figure S2). In most extreme cases, entire layers of cells were stacked, and while pure fluorescence emission is unchanged in such 3D configurations, the light captured in brightfield images takes a very different path compared to a monolayer (Figure S4). When the nucleus was detected, but the boundaries did not match, the possible causes were a different focal plane resulting in a shift, or mitosis that splits the stained nuclear DNA into two objects (Figure S6).

### Computational considerations of brightfield segmentation

Speed, memory consumption, and data requirements for training are important practical considerations. All tested models had tens of thousands of parameters, and take up between 5Gib and 7Gib of memory (15-175 Mib on disk). Segmenting an image with size 1080×1080 pixels takes 0.28 seconds for U-net and 6.64 seconds for Mask-RCNN, while the patching approach required by DeepCell renders its application impractical, with 159 seconds per image using a GPU.

Substantial amounts of data are required to train deep neural networks. We first tested models trained on data from a single cell line at a time. On average, we achieved accuracy within 6% of overall optimum with 32 full micrographs for each line, with moderate variability between the training sets (Figure 3A, B). This suggests that extensive manual labeling is required to train a new brightfield segmentation model.

**Figure 3.**
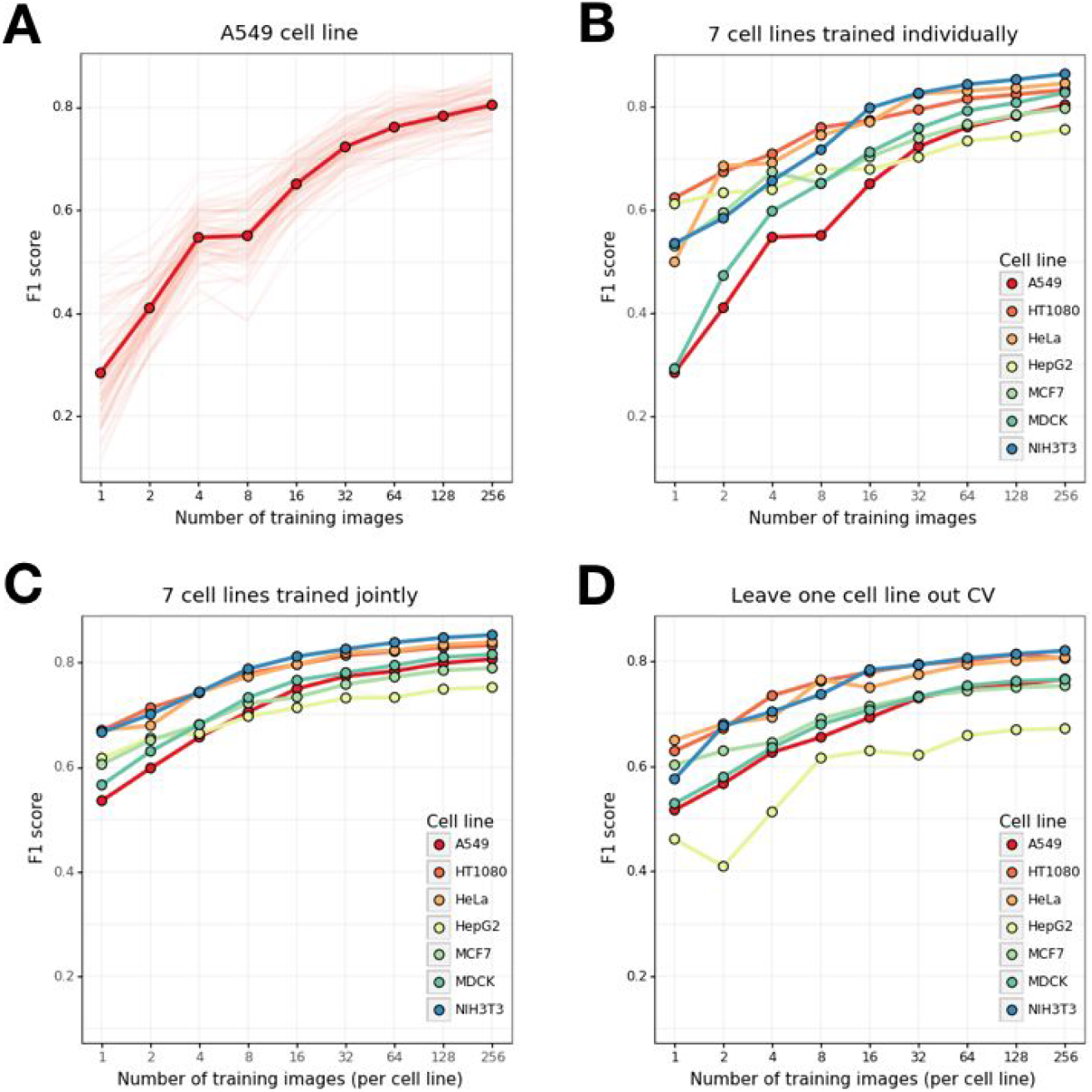
Segmentation performance depends on training set size. Pixel-level segmentation F1 score (y-axis) for increasing number of training images per cell line (x-axis). **A**. U-Net trained on A549 cell line only, with multiple random samples of training images (light lines), and their average (dark line and markers). **B**. As (A), but averages only for models trained separately on the seven different cell lines. **C**. As (B), but for models trained on a training set that includes the same fixed number of images from each cell line. **D**. As (C), but without including the tested cell line in the training set. Note that the total number of training images is seven times larger than x-axis value in (C), and six times larger than x-axis value in (D).

Next, we tested a model trained on many cell lines. Again, we varied the number of training images, and evaluated on held out data. The performance on any single line was improved by including the same number of images from other cell lines during training (Figure 3B,C), even when hundreds of training images were available for the target line. Conversely, removing images corresponding to the target line from a large diverse training set decreased performance on that line (Figure S7), suggesting that the trained networks do not generalize to previously unseen cell lines. However, if only a small number of up to eight training images was available to train a model for a line, an alternative model trained on the same number of images from each of the other six lines outperformed it (Figure 3B, 3D, S7). These results indicate that a pool of several annotated images or a large diverse training set is required for optimal performance.

### Data from multiple focal planes improves segmentation

Alternative focal planes contain different information about the sample, and it is not obvious which one is best to use for segmentation. Further, multiple acquisitions are feasible, as the brightfield images are quick to capture, and do not damage the specimen. Therefore, we next generated a dataset of human embryonic kidney cell (LNCaP) images with nine different focal planes in the brightfield channel (Figure 4A).

**Figure 4.**
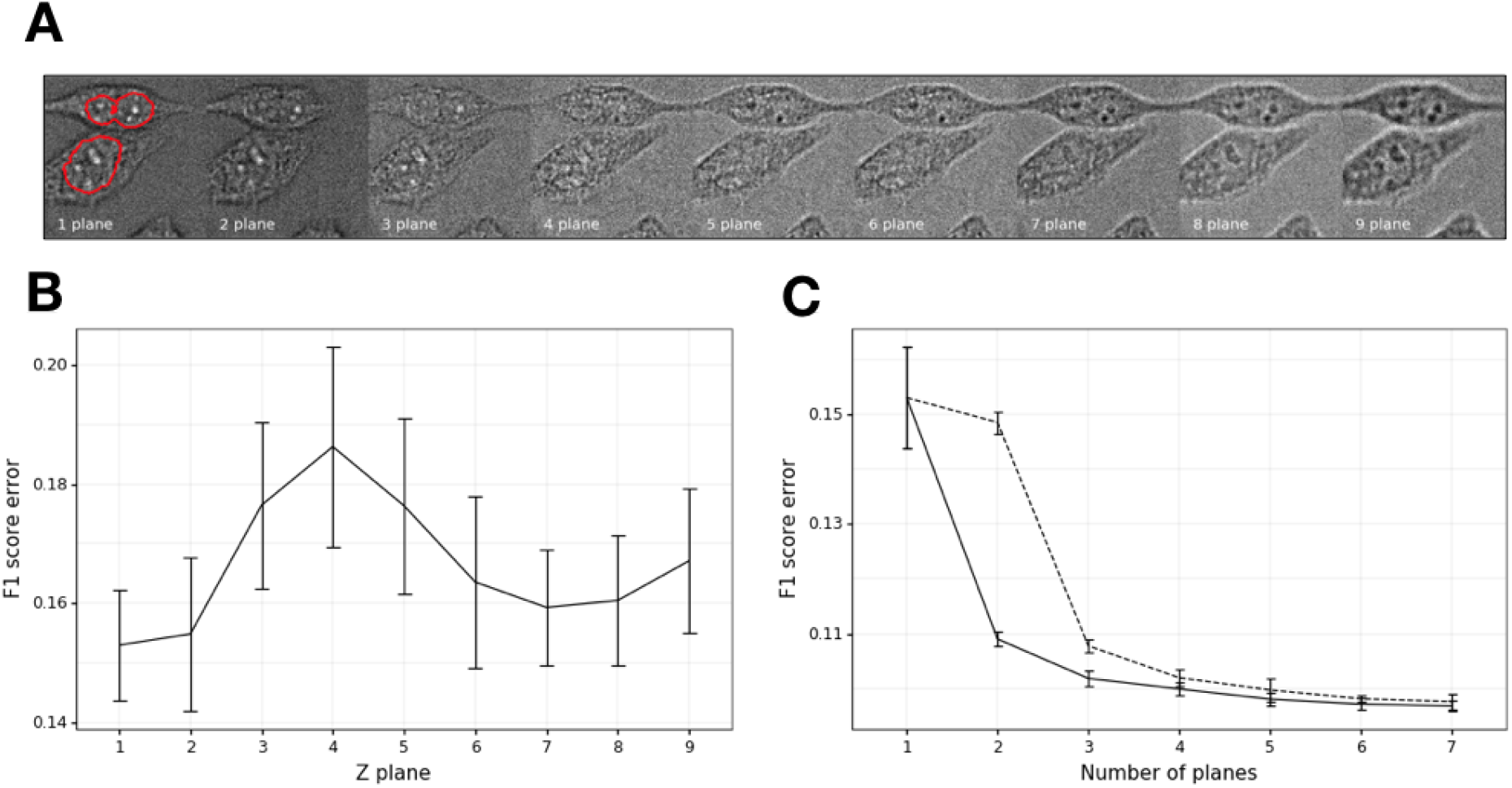
Multiple brightfield planes improve segmentation. **A**. Example brightfield acquisitions of the same specimen from nine focal planes, ranging from top (left) to bottom (right) in 1 micrometer steps. Red line: contour of the nucleus, as segmented from the fluorescence channel. **B**. F1 score error depends on the focal plane. Average error of four random training restarts (y-axis) for U-Net models trained on data from different focal planes (x-axis). Error bars – range of results. **C**. Two focal planes are sufficient. Average F1 score error across random plane selections (y-axis) for different number of input planes (x-axis) that are either all distinct (solid line), or repeat one of the input planes multiple times to control for the number of parameters (dashed line). Error bars – range of results.

First, we tested whether segmentation performance depends on the focal plane. We trained U-Net on all available data on each acquired plane, and evaluated on held out data. All planes had sufficient information to identify nuclei, but the central ones (planes 3 to 5) had a higher F1 error compared to the rest (0.18 vs 0.16, Figure 4B). One possible reason for this is the better contrast of subnuclear structures, such as nucleoli, in other planes (Figure 4A). Next, we considered whether using multiple input planes improves performance. We augmented the input layer of U-Net to accept a higher-dimensional input, leaving the rest of the architecture unchanged. Two layers improved accuracy over a single one (Figure 4C), while including information from additional ones gave diminishing returns.

## Discussion

We have demonstrated that multiple different neural network architectures can be trained to accurately segment nuclei from images of diverse cell lines acquired in fluorescence and brightfield channels. It is beneficial to have brightfield data from multiple focal planes, and annotations for dozens of training images to achieve optimal performance. Large training datasets from multiple different cell lines lead to more robust performance in segmenting nuclei. They help both in cases when only a few annotated images are available for a new cell line, but also when data are plentiful. This suggests that online batch learning will increase model performance over time.

Nearly all the pixel assignment errors can be ascribed a small number of interpretable causes. Many of these are due to fundamental discrepancies in ground truth data from fluorescence. For example, cells undergoing mitosis will identify with two ground truth nuclei from the Hoechst DNA stain, but as a single nucleus from a brightfield image. A slightly different focal plane, or switching imaging modes and returning to the same field of view can generate a shift between the two acquisition modes. Overlapping cells may also manifest differently in the two channels. However, as real-world applications evaluate cell population behavior, small segmentation discrepancies are often negligible, due to their minor contribution to the overall experimental variation.

We focused on the U-Net model for most of our investigation as the most practical approach. In our tests, it was about 25 times faster than Mask R-CNN, and about 500 times faster than DeepCell, making it usable in an online mode along with data acquisition. Its output is a smooth probability map that can be used as input to other methods for further refinement. This way, it does not suffer from the sharp thresholds of Mask R-CNN. Perhaps notably, the training process was subjectively easier using this architecture. The models trained faster, and were less fragile, with small changes having little effect on the performance.

While we could achieve good performance using U-net, many labeled images are required. A practical approach to using the network is to stain nuclei during the assay setup phase, and use the acquired images to perform the much simpler task of fluorescence-based segmentation. This annotation can then be used to generate a ground truth dataset that would serve to train or fine-tune a U-Net model for application on the experiment at scale.

## Methods

### Data collection

#### Seven cell lines data set

Seven different cell lines (mouse fibroblasts (NIH/3T3), canine kidney epithelial cells (MDCK), human cervical adenocarcinoma (HeLa), human breast adenocarcinoma (MCF7), human lung carcinoma (A549), human hepatocellular carcinoma (HepG2) and human fibrosarcoma (HT1080)) were seeded each into 48 wells of a CellCarrier-384 Ultra microplate (PerkinElmer, #6057300). The following cell numbers were seeded per well and cell line – NIH/3T3: 7,000, MDCK: 7,000, HeLa: 5,000, MCF7: 10,000, A549: 12,000, HepG2: 20,000, HT1080: 10,000. The following day the cells were fixed using 3.7% formaldehyde solution (Sigma Aldrich, #252549) and nuclei stained with 10μg/ml Hoechst 33342 (Thermo Fisher, #H3570). Brightfield and fluorescent images were acquired on an Opera Phenix™ high-content screening system (PerkinElmer) using a 20x water immersion objective in both non-confocal and confocal mode to capture fluorescence and brightfield images. 432 fields of view were captured for each cell line, for a total of 3024 images.

#### Nine focal planes data set

The commercially LNCaP cell line was grown in standard medium, plated to 96-well imaging plate (Cell Carrier Ultra, Perkin Elmer), fixed using formaldehyde, stained using DRAQ5 fluor (Abcam) to label nuclear DNA, and imaged on the CellVoyager 7000 (Yokogawa) using the 20x objective in both confocal mode to capture fluorescence images, and brightfield mode, in 9 z-planes using 1 micrometer steps. A total of 784 fields of view were captured, each at 9 focal heights.

#### Ground truth generation

Ground truth nuclei labels were created by Harmony, PerkinElmer proprietary image analysis software. Nuclei Detection method C was chosen for robustness with respect to size and fluorescence signal contrast variation of nuclei. The method uses local thresholding, and methods to split stuck nuclei combined with morphological opening, closing and filling. To improve ground truth quality we applied background flatfield correction to the fluorescent image (Kask et al. 2016). We further applied object filtering to exclude irregular shaped nuclei from consideration as training data. The segmentation quality was inspected manually across different cell lines and measurements.

### Training process

#### U-net model and training

The U-Net architecture used was based on (Ronneberger, Fischer, and Brox 2015), and consisted of contracting and an expansive paths of 1.3 million neurons in total. Contracting part was built using a series of three 3×3 convolutional filters, followed by rectified linear unit (ReLU) activation and 2×2 max pooling operation. Each step in the expansive path contained an up-sampling operation, series of convolutional layers followed by ReLU and concatenations with dimension-matched outputs from contracting layers. Adam optimization algorithm (Kingma and Ba 2014) was used with a learning rate of 0.0002 for updating network weights with respect to binary cross entropy loss. Loss was calculated on validation images after each training epoch (629 iterations) that consisted of 200 forward passes of 16 inputs of 288×288 pixels. Maximum number of epochs was 200, from which the model with lowest loss on validation images was selected. Learning rate was decreased by a factor of 10 when validation loss stopped improving for 10 consecutive epochs.

#### Mask-RCNN model and training

The Mask R-CNN architecture, was repurposed from (He et al. 2018), and updated to accept 512×512 pixel images as input. Stochastic gradient descent was used with a learning rate of 0.001. Loss was calculated on validation images after each training epoch of 600 inputs with a batch size of 1. Maximum number of epochs was set to 130, from which the model with lowest loss on validation images was selected. Learning rate was decreased by a factor of 5 when validation loss stopped improving for 10 consecutive epochs.

#### DeepCell model and training

The DeepCell architecture (Van Valen et al. 2016) consisted of four layers of 64 convolutional filters with ReLU activation, each followed by 2×2 max pooling, and a deeply connected layer. Adam optimization algorithm was used with a learning rate of 0.001 with respect to categorical cross-entropy loss. Loss was calculated on 31×31 pixel patches from validation images after each training epoch that consisted of 249,654 inputs with a batch size of 4096. The model was trained for 200 epochs from which the best model with respect to validation loss was picked.

#### Training datasets

The 3024 images from seven cell lines were divided into three independent sets: training set with 2016 images, validation set with 504 images, and test set with 504 images, with different representations from the seven lines (Table S1). The nine focal planes dataset was similarly split once into 628 training, 78 validation, and 78 test images. All training and evaluation took place on this single split of images for both datasets.

#### Training on individual cell lines

For each of the seven cell lines, we constructed nine training sets (1, 2, 4, 8, 16, 32, 64, 128, and 256 images; 7×9 = 63 datasets total). We trained a U-Net model on each. Resulted models were validated and tested using cell line specific validation and test sets.

#### Training on all cell lines jointly

We constructed nine training sets, each containing a fixed number of images from each of the seven cell lines (1, 2, 4, 8, 16, 32, 64, 128, and 256 images per cell line; dataset sizes 7, 14, 28, 56, 112, 224, 448, 896, 1792 images). We trained a U-Net model on each, and validated on a joint validation set.

#### Leave-one-out training

For each cell line, we constructed nine training sets, each containing a fixed number of images from each of the other six cell lines (1, 2, 4, 8, 16, 32, 64, 128, and 256 images per cell line; dataset sizes 6, 12, 24, 48, 96, 192, 384, 768, 1536 images). We trained a U-Net model on each, and validated on the target cell line specific validation set.

#### Multi-plane data – single plane training

The 784 images were divided into three independent sets: training set with 628 images, validation set with 78 images, and test set with 78 images. The rest of the training was performed as for seven cell lines data.

#### Multi-plane data – multi-plane training

For each number of *k* input layers, we augmented the U-Net structure to accept *k* input channels, with *k* ranging from 1 to 9. We then trained the network as before, both using *k* planes randomly sampled without replacement, as well as *k-1* planes randomly sampled without replacement, and one plane repeated twice. This allows for a direct evaluation of the influence of adding an extra input channel, as the number of model parameters does not change.

## Author contributions

Performed model comparisons: DF, SOS. Performed model evaluation on multiple cell lines: DF, SOS. Performed model evaluation on multiple focal planes: DF, DM. Generated training data: SP, JW, AS, KP. Supervised project: KP, LP. Wrote paper: DF, SOS, LP.

Grant support: IUT34-4 from Estonian Research Council and Wellcome to LP.

## Supporting information

**Figure S1.**
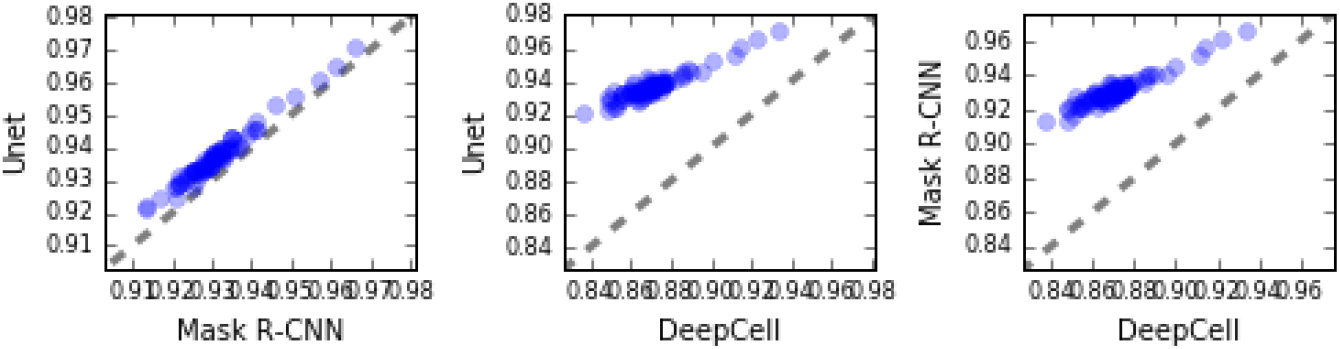
Comparison of per-image test accuracy. Per-image accuracy (x and y axis, blue markers) for pairs of tested models (U-Net, Mask-RCNN and DeepCell). Dashed line: y=x.

**Figure S2.**
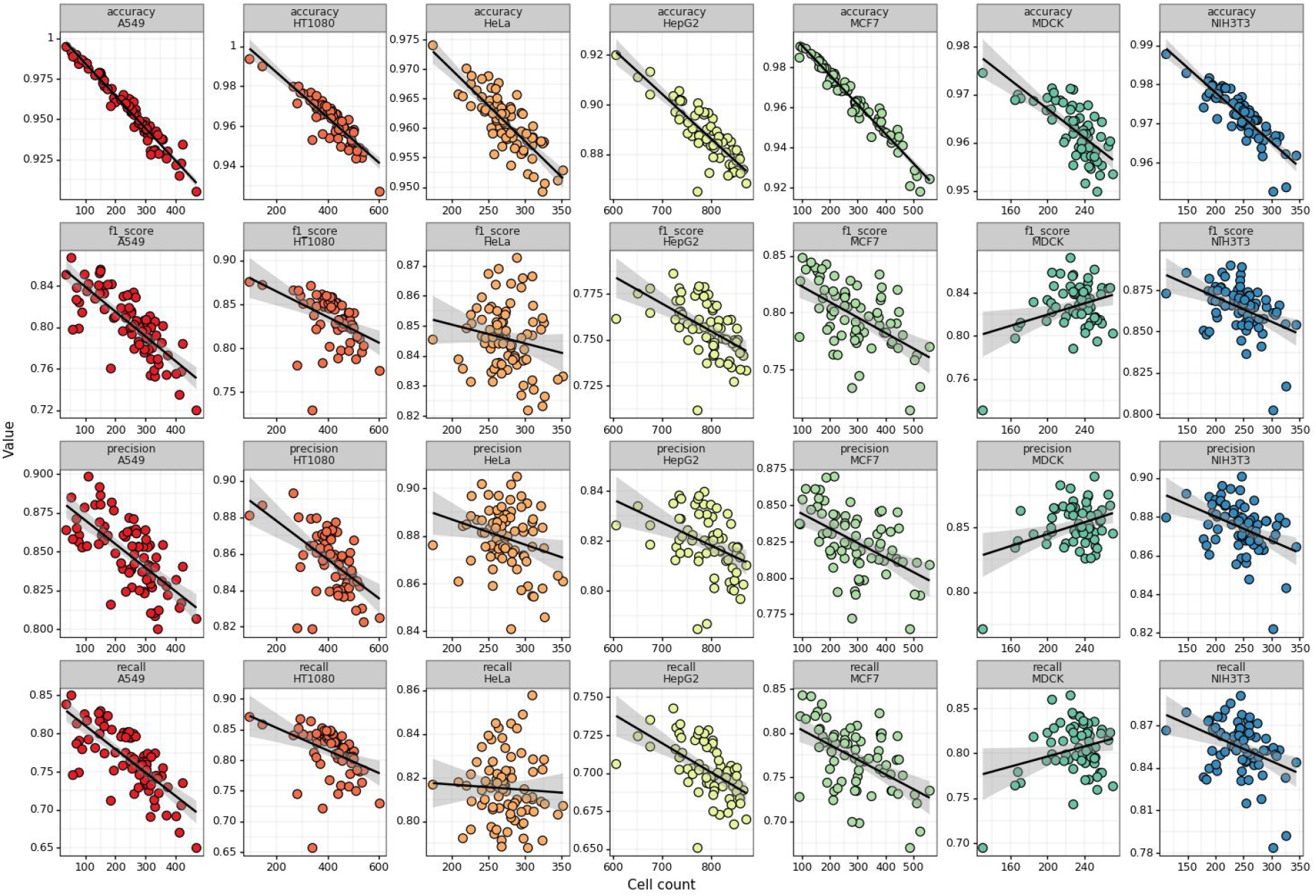
Per-image accuracy metrics depend on the number of cells in the image. Cell count (x-axis) contrasted against a performance metric (y-axis) for each of the seven cell lines (columns), and using different metrics (first row: accuracy; second row: F1 score; third row: precision; fourth row: recall). Solid line: linear regression fit; shaded area: 95% confidence interval.

**Figure S3.**
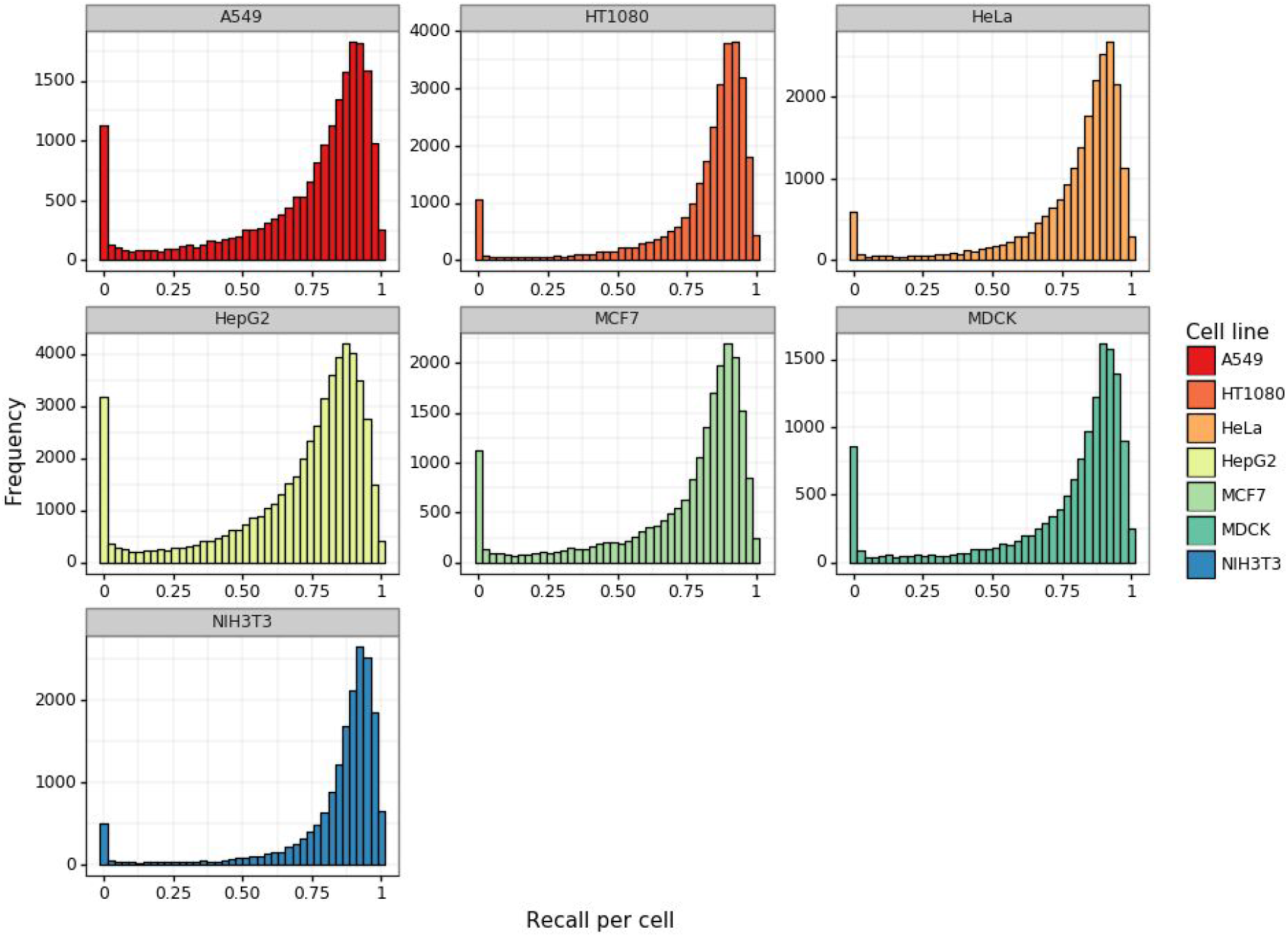
Distribution of per-cell recall. Frequency (y-axis) of per-cell recall (x-axis) for each of the seven cell lines (panels).

**Figure S4.**
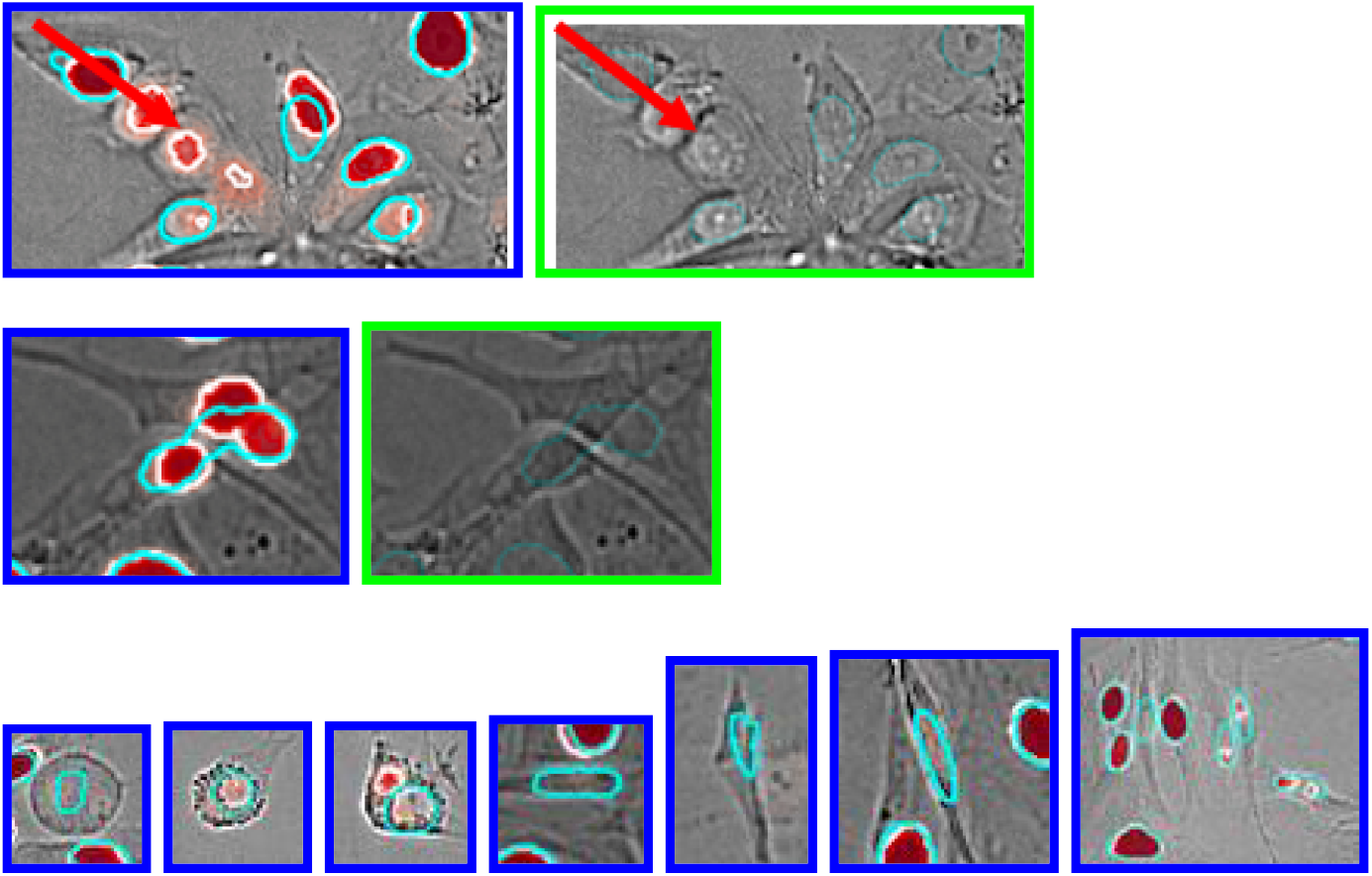
Examples of cells with unusual morphology, and dense stacking (inaccurate nuclei calls). Teal outline: true positive nucleus boundary. Red: areas of high posterior probability. Red arrow: highlighted cell. Blue image border: brightfield, ground truth, and posterior prediction. Green image border: brightfield and ground truth only matching the annotated image on the left.

**Figure S5.**
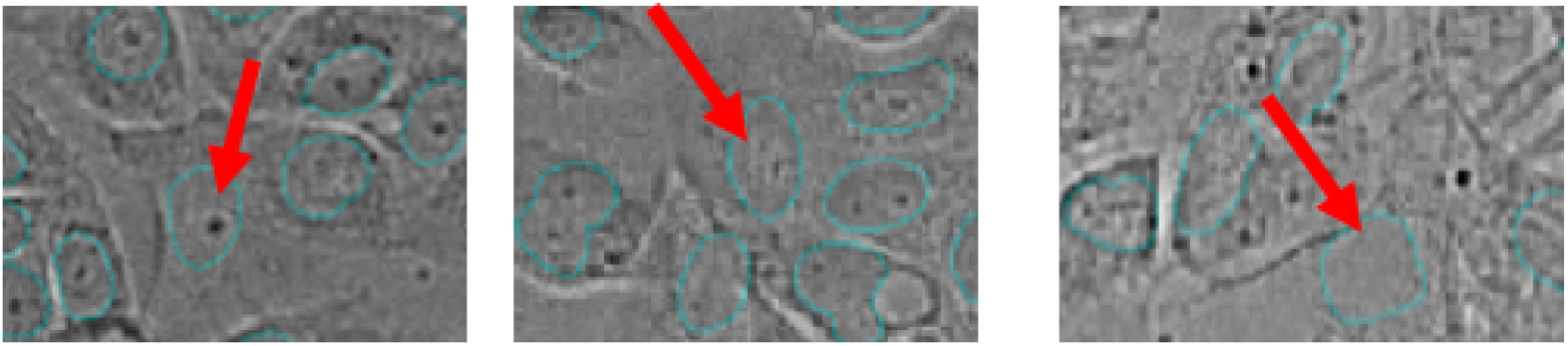
Examples of cells with low signal in brightfield channel (false negative nuclei calls). Teal outline: true positive nucleus boundaries. Red arrows: true positive nuclei with low posterior probability (probability not shown).

**Figure S6.**
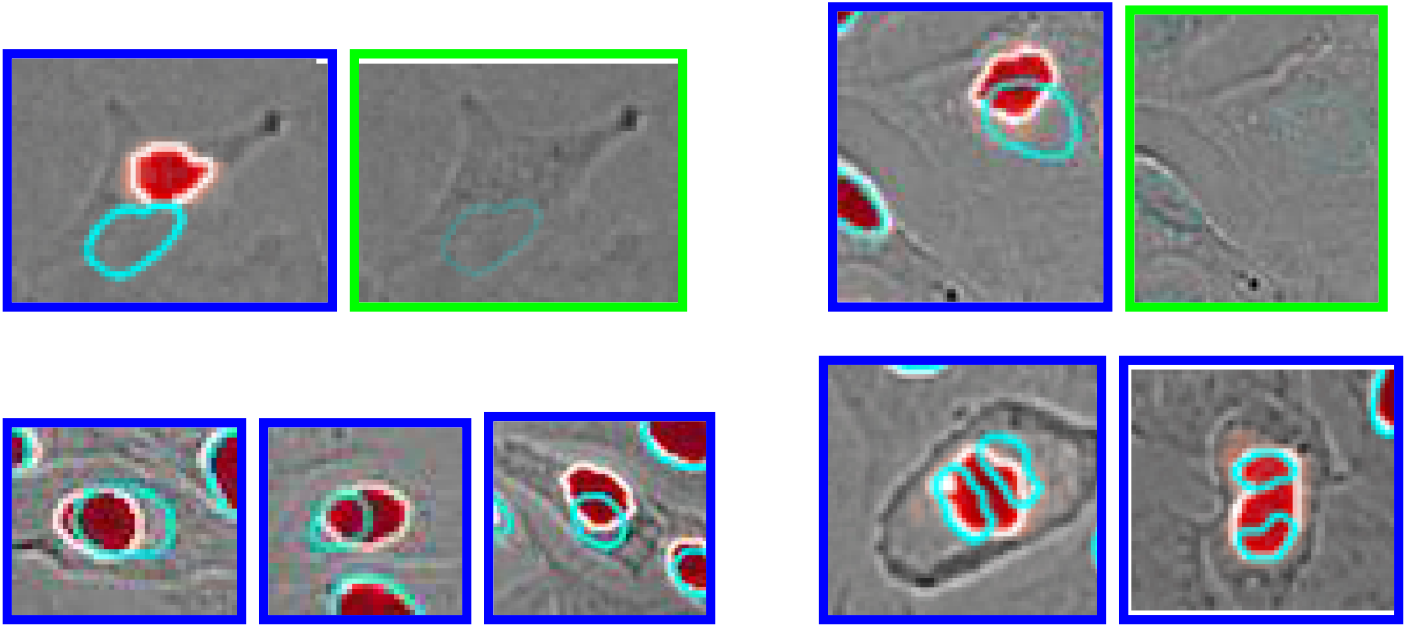
Examples of cells likely shifted in nuclear location between focal planes, or undergoing mitosis (false positive nuclear region calls). Teal outline: true positive nucleus boundary. Red: areas of high posterior probability. Red arrow: highlighted cell. Blue image border: brightfield, ground truth, and posterior prediction. Green image border: brightfield and ground truth only.

**Figure S7.**
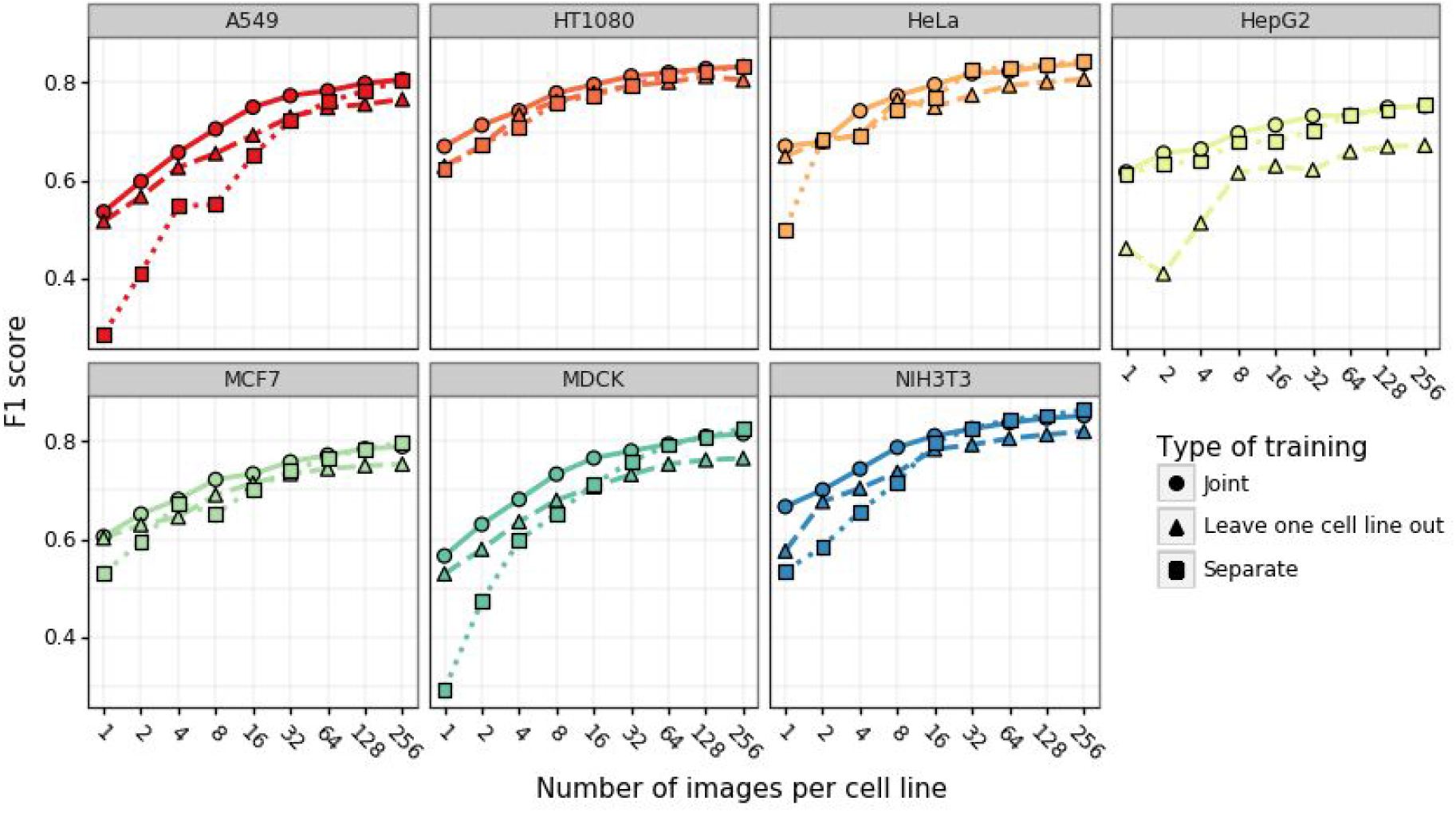
Effect of training dataset size and composition on model performance. F1 score on held-out data (y-axis) for increasing number of training images (x-axis) for each of seven cell lines (panels), training either on the target line only (squares, dotted line), all lines together (circles, solid line), or on all the lines but the target one (triangles, dashed line).

**Supplementary Table 1:** Distribution of images in seven cell lines data set.

|  | **Train** | **Validation** | **Test** |
| --- | --- | --- | --- |
| **A549** | 286 | 66 | 80 |
| **HT1080** | 284 | 78 | 70 |
| **HeLa** | 293 | 58 | 81 |
| **HepG2** | 283 | 82 | 67 |
| **MCF7** | 290 | 70 | 72 |
| **MDCK** | 292 | 79 | 61 |
| **NIH3T3** | 288 | 71 | 73 |
| **Total** | **2016** | **504** | **504** |

**Supplementary Table 2:** Performance of the U-Net jointly trained on brightfield images from all cell lines. Table entries for different cell lines (rows) are the errors according to the metric specified in the column, i.e. 1-metric.

| **Cell line** | **Accuracy** | **F1-score** | **Precision** | **Recall** | **Specificity** |
| --- | --- | --- | --- | --- | --- |
| **A549** | 0.0429 | 0.1829 | 0.1571 | 0.2065 | 0.0203 |
| **HT1080** | 0.0367 | 0.1618 | 0.1462 | 0.1765 | 0.0184 |
| **HeLa** | 0.0392 | 0.1544 | 0.1232 | 0.1832 | 0.0175 |
| **HepG2** | 0.1111 | 0.2397 | 0.1857 | 0.2868 | 0.0535 |
| **MCF7** | 0.0369 | 0.2024 | 0.1668 | 0.2345 | 0.0162 |
| **MDCK** | 0.0378 | 0.1734 | 0.1539 | 0.1917 | 0.0185 |
| **NIH3T3** | 0.0288 | 0.1394 | 0.1387 | 0.1399 | 0.0161 |

**Supplementary Table 3:** Performance of the U-Net trained on brightfield images from each cell line separately. Table entries for different cell lines (rows) are the errors according to the metric specified in the column, i.e. 1-metric.

| **Cell line** | **Accuracy** | **F1-score** | **Precision** | **Recall** | **Specificity** |
| --- | --- | --- | --- | --- | --- |
| **A549** | 0.0429 | 0.1855 | 0.1511 | 0.2165 | 0.0191 |
| **HT1080** | 0.0364 | 0.1609 | 0.1436 | 0.1770 | 0.0180 |
| **HeLa** | 0.0378 | 0.1487 | 0.1186 | 0.1766 | 0.0169 |
| **HepG2** | 0.1094 | 0.2364 | 0.1807 | 0.2849 | 0.0518 |
| **MCF7** | 0.0363 | 0.1954 | 0.1749 | 0.2144 | 0.0177 |
| **MDCK** | 0.0355 | 0.1622 | 0.1446 | 0.1788 | 0.0175 |
| **NIH3T3** | 0.0265 | 0.1288 | 0.1237 | 0.1337 | 0.0142 |

**Supplementary Table 4:** Performance of the U-Net trained on fluorescent images from all cell lines jointly. Table entries for different cell lines (rows) are the errors according to the metric specified in the column, i.e. 1-metric.

| **Cell line** | **Accuracy** | **F1-score** | **Precision** | **Recall** | **Specificity** |
| --- | --- | --- | --- | --- | --- |
| **A549** | 0.0068 | 0.0285 | 0.0157 | 0.0408 | 0.0022 |
| **HT1080** | 0.0035 | 0.0152 | 0.0154 | 0.0150 | 0.0020 |
| **HeLa** | 0.0126 | 0.0500 | 0.0153 | 0.0821 | 0.0022 |
| **HepG2** | 0.0164 | 0.0335 | 0.0189 | 0.0477 | 0.0060 |
| **MCF7** | 0.0034 | 0.0178 | 0.0211 | 0.0144 | 0.0023 |
| **MDCK** | 0.0037 | 0.0171 | 0.0156 | 0.0184 | 0.0020 |
| **NIH3T3** | 0.0021 | 0.0102 | 0.0115 | 0.0088 | 0.0014 |

**Supplementary Table 5:** Performance of the U-Net model trained on fluorescent images from each cell line separately. Table entries for different cell lines (rows) are the errors according to the metric specified in the column, i.e. 1-metric.

| **Cell line** | **Accuracy** | **F1-score** | **Precision** | **Recall** | **Specificity** |
| --- | --- | --- | --- | --- | --- |
| **A549** | 0.0072 | 0.0303 | 0.0146 | 0.0454 | 0.0020 |
| **HT1080** | 0.0036 | 0.0156 | 0.0169 | 0.0143 | 0.0022 |
| **HeLa** | 0.0158 | 0.0635 | 0.0140 | 0.1081 | 0.0019 |
| **HepG2** | 0.0153 | 0.0313 | 0.0182 | 0.0439 | 0.0058 |
| **MCF7** | 0.0036 | 0.0189 | 0.0213 | 0.0163 | 0.0023 |
| **MDCK** | 0.0041 | 0.0185 | 0.0238 | 0.0131 | 0.0030 |
| **NIH3T3** | 0.0025 | 0.0118 | 0.0183 | 0.0052 | 0.0022 |

## Bibliography

Altmann, Richard. 1894. Die Elementarorganismen Und Ihre Beziehungen Zu Den Zellen.

Angermueller, Christof, Tanel Pärnamaa, Leopold Parts, and Oliver Stegle. 2016. “Deep Learning for Computational Biology.” Molecular Systems Biology 12 (7): 878.

Bray, Mark-Anthony, Shantanu Singh, Han Han, Chadwick T. Davis, Blake Borgeson, Cathy Hartland, Maria Kost-Alimova, Sigrun M. Gustafsdottir, Christopher C. Gibson, and Anne E. Carpenter. 2016. “Cell Painting, a High-Content Image-Based Assay for Morphological Profiling Using Multiplexed Fluorescent Dyes.” Nature Protocols 11 (9): 1757–74.

Brown, Robert. 1833. “XXXV. On the Organs and Mode of Fecundation in Orchideae and Asclepiadeae.” Transactions of the Linnean Society of London. https://doi.org/10.1111/j.1095-8339.1829.tb00158.x.

Christiansen, Eric M., Samuel J. Yang, D. Michael Ando, Ashkan Javaherian, Gaia Skibinski, Scott Lipnick, Elliot Mount, et al. 2018. “In Silico Labeling: Predicting Fluorescent Labels in Unlabeled Images.” Cell 173 (3): 792–803.e19.

Falk, Thorsten, Dominic Mai, Robert Bensch, Özgün Çiçek, Ahmed Abdulkadir, Yassine Marrakchi, Anton Böhm, et al. 2019. “U-Net: Deep Learning for Cell Counting, Detection, and Morphometry.” Nature Methods 16 (1): 67–70.

Fan, Haoqiang, and Erjin Zhou. 2016. “Approaching Human Level Facial Landmark Localization by Deep Learning.” Image and Vision Computing. https://doi.org/10.1016/j.imavis.2015.11.004.

He, Kaiming, Georgia Gkioxari, Piotr Dollar, and Ross Girshick. 2018. “Mask R-CNN.” IEEE Transactions on Pattern Analysis and Machine Intelligence, June. https://doi.org/10.1109/TPAMI.2018.2844175.

Hooke, Robert, and Jo Martyn And. 1665. “Micrographia, Or, Some Physiological Descriptions of Minute Bodies Made by Magnifying Glasses :with Observations and Inquiries Thereupon /by R. Hooke.” https://doi.org/10.5962/bhl.title.904.

Isherwood, Beverley, Paul Timpson, Ewan J. McGhee, Kurt I. Anderson, Marta Canel, Alan Serrels, Valerie G. Brunton, and Neil O. Carragher. 2011. “Live Cell in Vitro and in Vivo Imaging Applications: Accelerating Drug Discovery.” Pharmaceutics 3 (2): 141–70.

Jones, William, Kaur Alasoo, Dmytro Fishman, and Leopold Parts. 2017. “Computational Biology: Deep Learning.” Emerging Topics in Life Sciences. https://doi.org/10.1042/etls20160025.

Kask, Peet, Kaupo Palo, Chris Hinnah, and Thora Pommerencke. 2016. “Flat Field Correction for High-Throughput Imaging of Fluorescent Samples.” Journal of Microscopy 263 (3): 328–40.

Kingma, Diederik P., and Jimmy Ba. 2014. “Adam: A Method for Stochastic Optimization.” http://arxiv.org/abs/1412.6980.

Li, Linfeng, Qiong Zhou, Ty C. Voss, Kevin L. Quick, and Daniel V. LaBarbera. 2016. “High-Throughput Imaging: Focusing in on Drug Discovery in 3D.” Methods 96 (March): 97–102.

Pärnamaa, Tanel, and Leopold Parts. 2017. “Accurate Classification of Protein Subcellular Localization from High-Throughput Microscopy Images Using Deep Learning.” G3 7 (5): 1385–92.

Rohban, Mohammad Hossein, Shantanu Singh, Xiaoyun Wu, Julia B. Berthet, Mark-Anthony Bray, Yashaswi Shrestha, Xaralabos Varelas, Jesse S. Boehm, and Anne E. Carpenter. 2017. “Systematic Morphological Profiling of Human Gene and Allele Function via Cell Painting.” eLife 6 (March). https://doi.org/10.7554/eLife.24060.

Ronneberger, Olaf, Philipp Fischer, and Thomas Brox. 2015. “U-Net: Convolutional Networks for Biomedical Image Segmentation.” Lecture Notes in Computer Science. https://doi.org/10.1007/978-3-319-24574-4_28.

Usaj, Mojca Mattiazzi, Erin B. Styles, Adrian J. Verster, Helena Friesen, Charles Boone, and Brenda J. Andrews. 2016. “High-Content Screening for Quantitative Cell Biology.” Trends in Cell Biology. https://doi.org/10.1016/j.tcb.2016.03.008.

Van Valen, David A., Takamasa Kudo, Keara M. Lane, Derek N. Macklin, Nicolas T. Quach, Mialy M. DeFelice, Inbal Maayan, Yu Tanouchi, Euan A. Ashley, and Markus W. Covert. 2016. “Deep Learning Automates the Quantitative Analysis of Individual Cells in Live-Cell Imaging Experiments.” PLOS Computational Biology. https://doi.org/10.1371/journal.pcbi.1005177.

